## Supplementary figures with legends for "Modeling corticotroph deficiency with pituitary organoids supports the functional role of *NFKB2* in human pituitary differentiation"

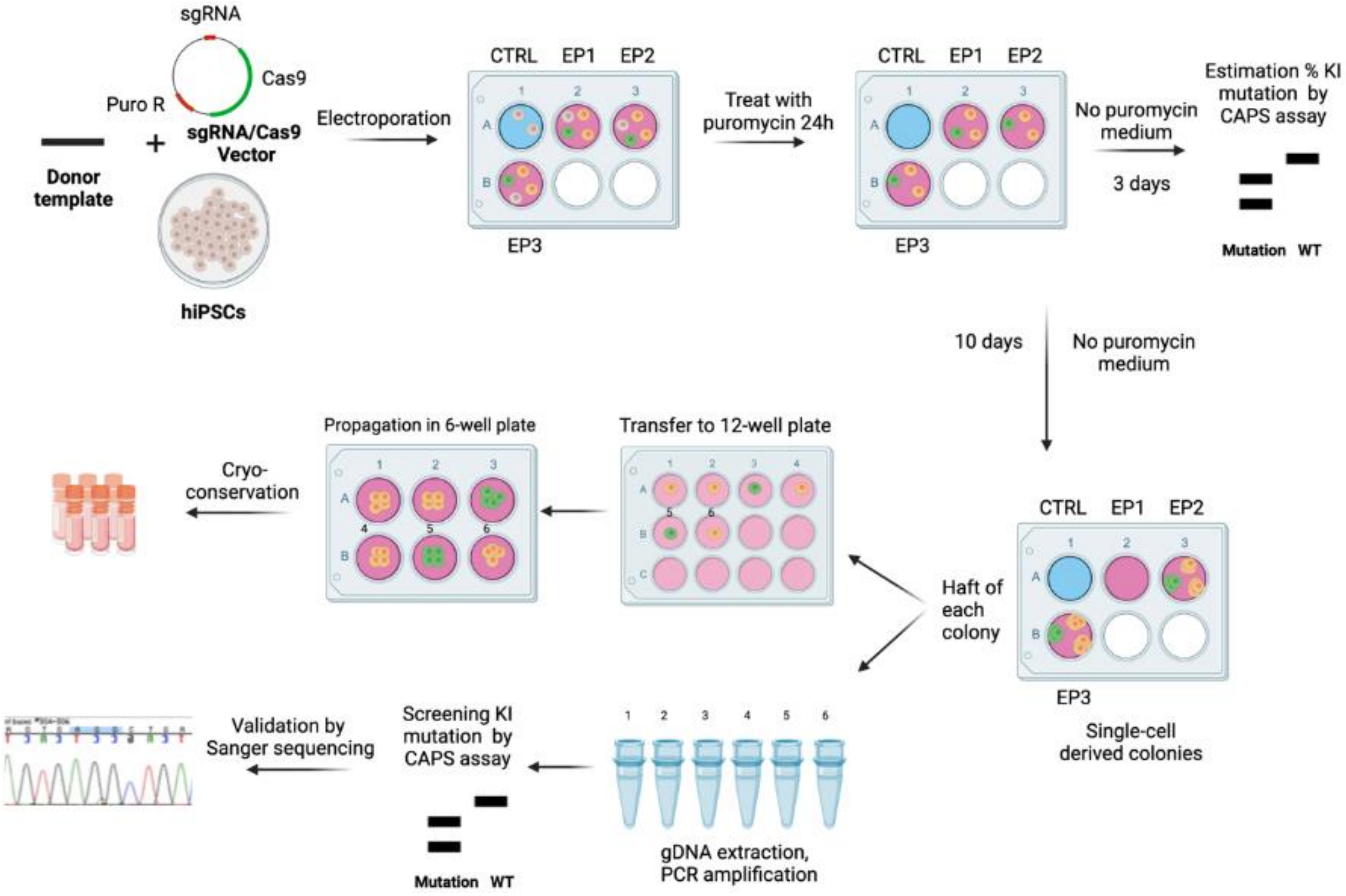


**Suppl. Figure S1: Summary of the key steps of CRISPR/Cas9-mediated genome knock-in editing in hiPSC.** The appropriate CRISPR/Cas9 vector and ssODN were introduced in hiPSC by electroporation. Resistant cells were selected by puromycin for 24 hours. Percentage of mutants was estimated by a CAPS assay. Mutant cells were amplified and manually picked after 10 days. Half of each clonal colony was used for screening and verified by Sanger sequencing. The other half was used for amplification and then stocked by cryo-conservation. Non-edited cells (in gray), edited cells without desired mutations (in yellow) and edited cells with desired mutation (in green) are depicted.

*sgRNA: single guide RNA; puro R: puromycin resistant sequence; CTRL: control; EP1, EP2, EP3: three wells containing hiPSC electroporated with the vectors and donor template; CAPS: cleaved amplified polymorphic sequences; WT: wild-type; KI: knock-in.*

A

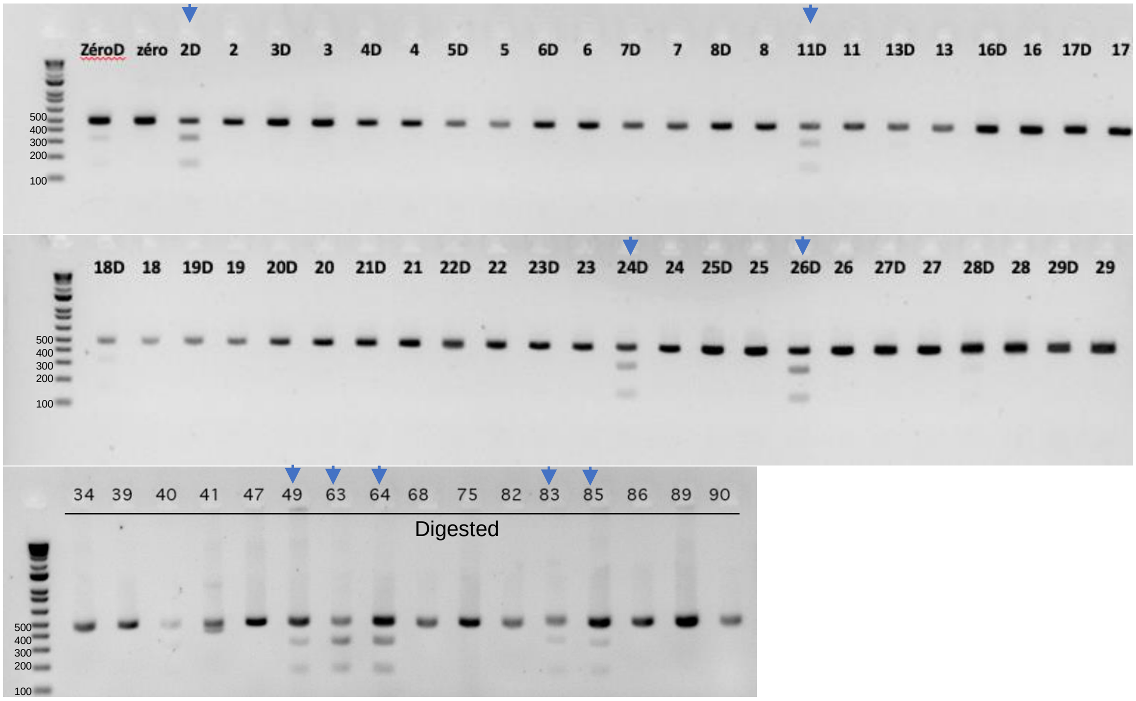


B

| Clone N° | Sanger sequencing |
|---|---|
| 2 | *TBX19*$^{K146R/fs}$ |
| 11 | *TBX19*$^{K146R/fs}$ |
| 24 | *TBX19*$^{K146R/fs}$ |
| 26 | *TBX19*$^{K146R/WT}$ |
| 49 | *TBX19*$^{fs/fs}$ |
| 63 | *TBX19*$^{K146R/K146R}$ |
| 64 | *TBX19*$^{fs/fs}$ |
| 83 | N/A |
| 85 | *TBX19*$^{fs/fs}$ |

**Suppl Figure S2: Results of CAPS assay and Sanger sequencing analysis for editing *TBX19* mutation**

**A**, CAPS assay with MseI restriction enzyme in hiPSC population (zero) on day 3 post-transtection and screening individual colony. WT (472 bp band); Blocking mutant digested (D, 2 bands at 314 bp and 158 bp). Positif clones which had 3 bands (marked by an blue arrow) were then confirmed by Sanger sequencing.

**B**, Sanger sequencing analysis of positives clones. There was one clone (# 63) homozygous *TBX19*$^{K146R/K146R}$ . There were heterozygous *TBX19*$^{K146R/WT}$ , *TBX19*$^{K146R/fs}$ and *TBX19*$^{fs/fs}$ . Frameshift (fs).

*N/A: not available*

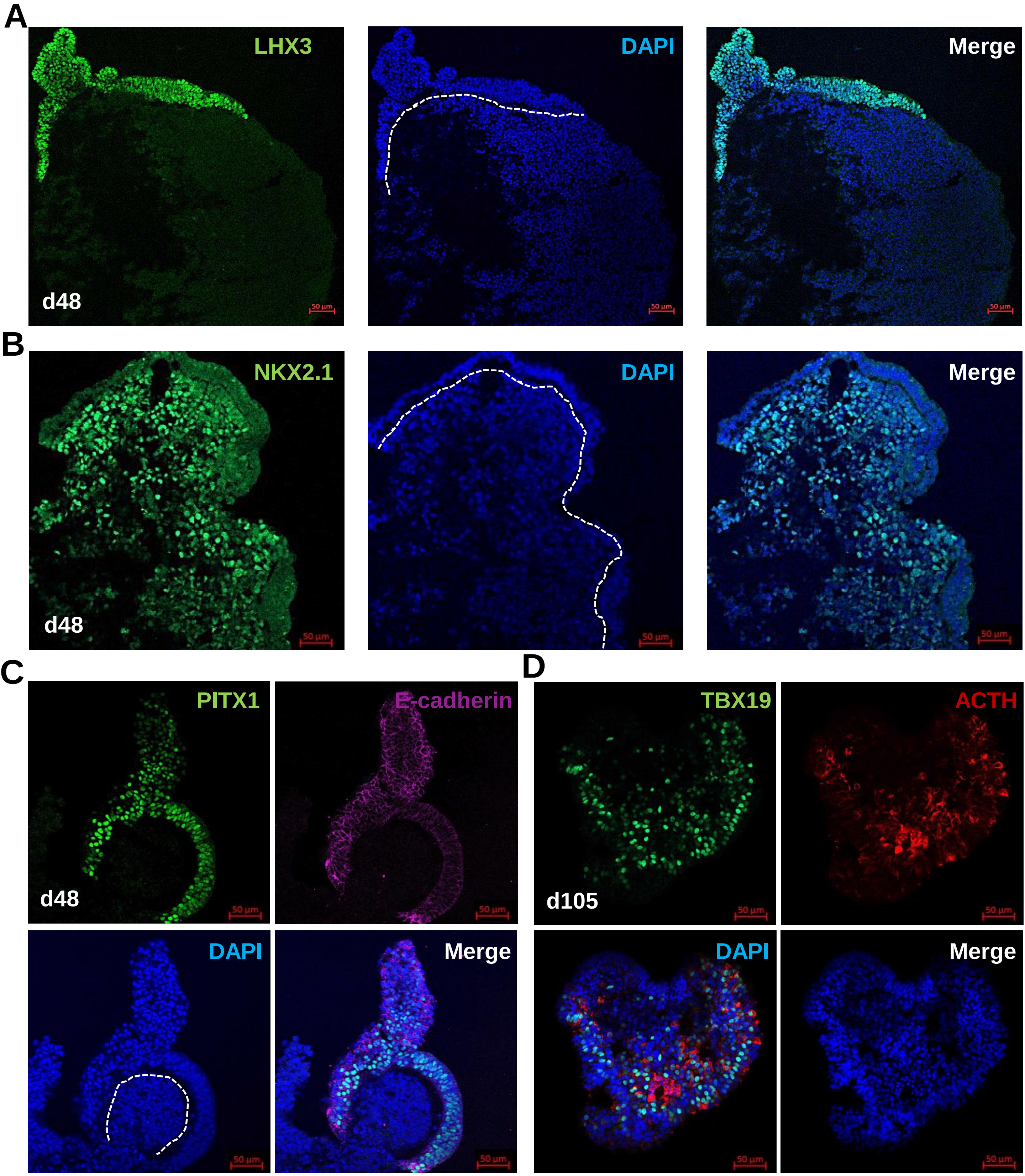


**Suppl Figure S3: Differentiation of hiPSC control into pituitary organoid using 3D culture**
**A-C**, Induction of two tissues hypothalamus-pituitary in organoid on day 48.
**A,** Oral ectoderm-like tissue expressed pituitary progenitor markers (LHX3).
**B,** Hypothalamus-like tissue expressed hypothalamic progenitor markers (NKX2.1).
**C,** Oral ectoderm-like tissue expressed PITX1 and E-cadherin.
**D,** Corticotroph cells expressed TBX19 in the nucleus and ACTH in the cytoplasm in organoid on day 105.
Scale bars: 50 µm

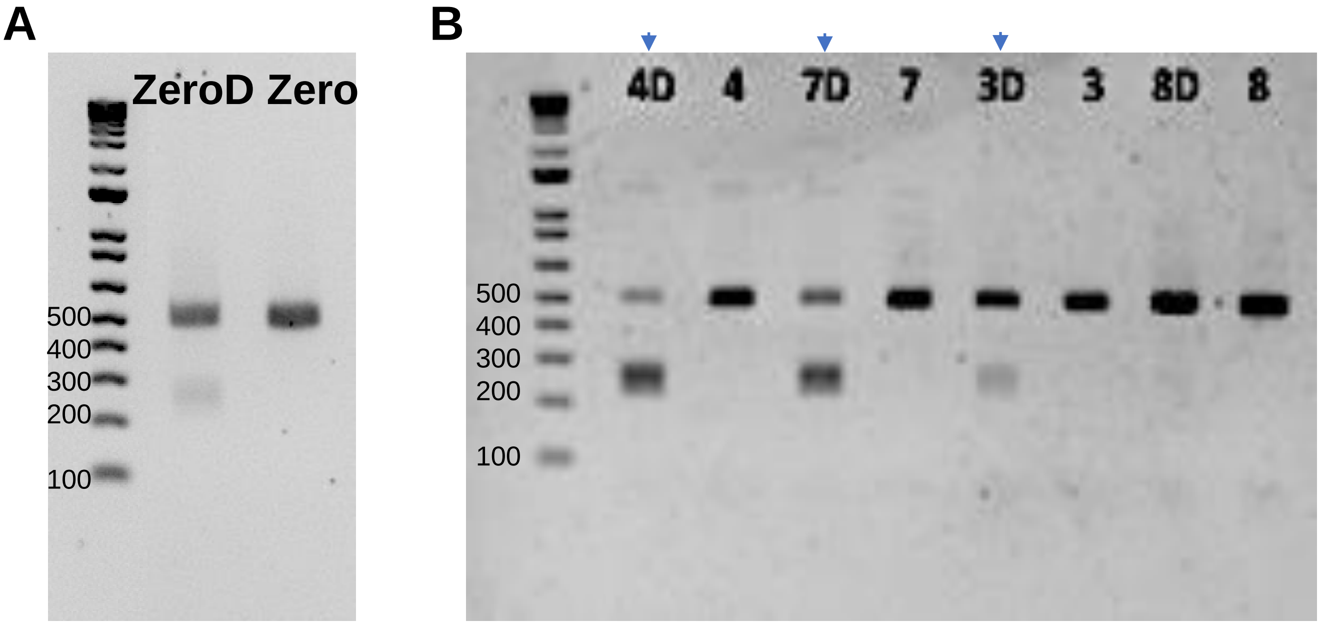


C

| Clone N° | Sanger sequencing |
|---|---|
| 3 | *NFKB2*$^{D865G/WT}$ |
| 4 | *NFKB2*$^{D865G/D865G}$ |
| 7 | *NFKB2*$^{D865G/D865G}$ |

**Suppl Figure S4: Results of CAPS assay and Sanger sequencing for editing *NFKB2* mutation**

**A**, CAPS assay with BtsI restriction enzyme in hiPSC population (zero) on day 3 post-transtection. WT or Undigested (498 bp band); Digested or mutant (D, 2 bands at 239 bp and 259 bp).

**B**, CAPS assay with BtsI restriction enzyme for screen individual colony. D: digested. Positif clones which had 3 bands (marked by an blue arrow) were then confirmed by Sanger sequencing.

**C**, Sanger sequencing analysis of positives clones. 2 clones (#4 and #7) were homozygous *NFKB2*$^{D865G/D865G}$ . One clone was heterozygous *NFKB2*$^{D865G/WT}$.

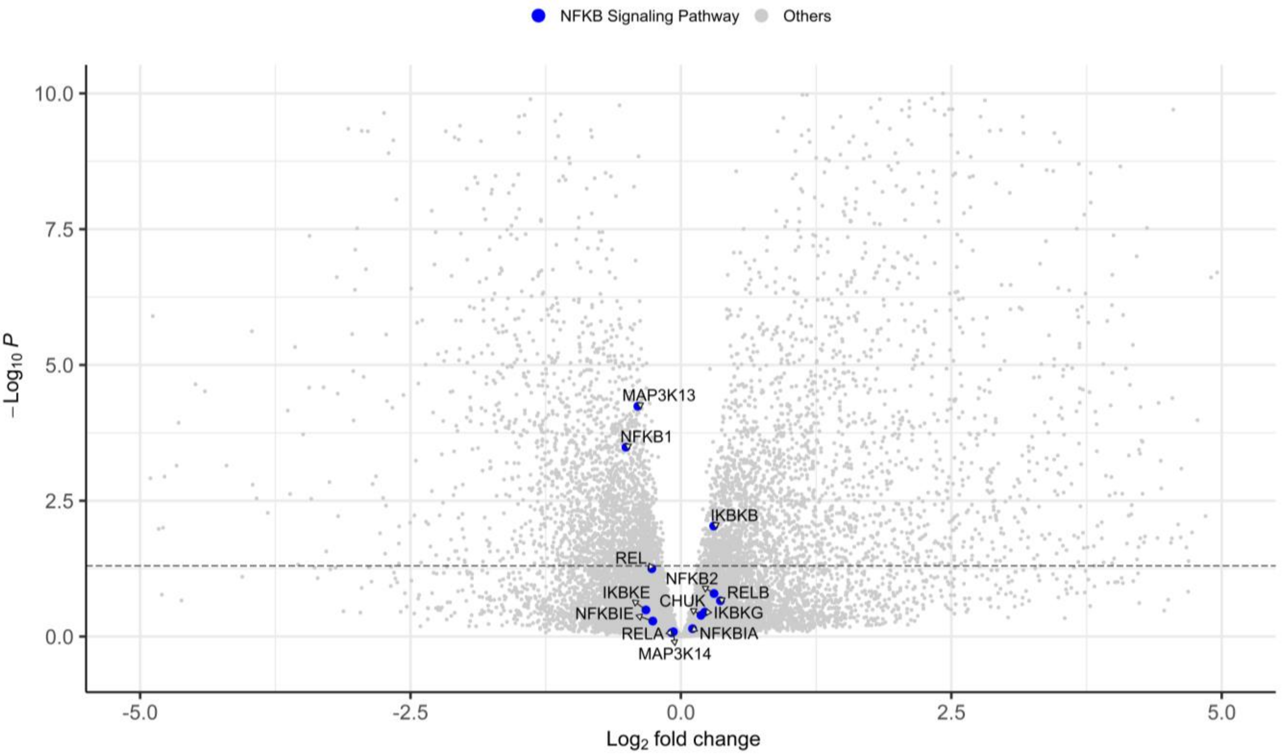


**Suppl Figure S5:** Volcano plot of genes showing differential gene expression between WT vs *NFKB2*$^{D865G/D865G}$ organoids for genes involved in NFKB canonical and non-canonical pathways

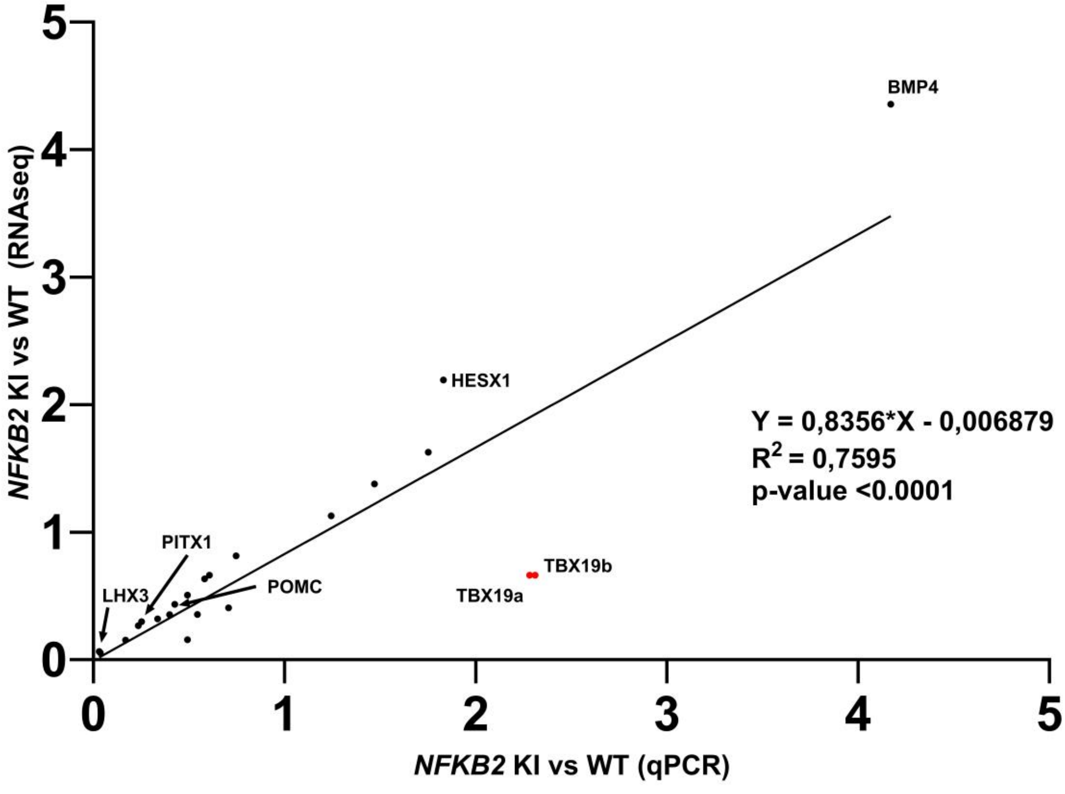


**Suppl Figure S6:** Correlation analysis of qRT-PCR and RNA-seq on same samples Close correlation was found between ratios measured by RNAseq and RT-qPCR when measured on same samples. Only transcripts levels measurement for *TBX19* (red dots) where not correlated. The 2 red dots represent two different sets of primers used for *TBX19* RT-qPCR
