## Supplementary data for "Modeling corticotroph deficiency with pituitary organoids supports the functional role of *NFKB2* in human pituitary differentiation"

**Supplemental data**

**Suppl. Tables**

| **Line** | **Sex** | **Age** | **Derivation** | **Reprogramming method** | **Reprogramming factors** |
| --- | --- | --- | --- | --- | --- |
| 10742L | Female | 82 | Fibroblast | Nucleofection | OCT4, SOX2, KLF4, c-MYC, SH-P53 |

**Table S1: hiPSC control line used in the study**

| **Nomenclature used in this paper** | **Sequence** |
| --- | --- |
| sgRNA | AAGCTGACCAACAAGCTCAA |
| sgRNA oligo up | 5’-CACCGAAGCTGACCAACAAGCTCAA-3’ |
| sgRNA oligo down | 5’-AAACTTGAGCTTGTTGGTCAGCTTC-3’ |
| ssODN | TGGATGAAAGCTCCCATCTCCTTCAG  CAAAGTGAGGCTGACCAACAAGTTaAA  TGGAGGCGGGCAGGTACGAATGAGG  CGGGCAGGCCTGGCCACCCGCT |

**Table S2: List of oligonucleotides used for CRISPR/Cas9 experiments to target *TBX19^K146R^***

| **Nomenclature used in this paper** | **Sequence** |
| --- | --- |
| sgRNA | GTGAAGGAAGACAGTGCGTA |
| sgRNA oligo up | 5’-CACCGTGAAGGAAGACAGTGCGTA-3’ |
| sgRNA oligo down | 5’-AAACTACGCACTGTCTTCCTTCAC-3’ |
| ssODN | TCCCATTCCTGTCCCCATTTACCCCC  AGCAGAGGTGAAGGAAGGCAGTGCC  TACGGGAGCCAGTCAGTGGAGCAGG  AGGCAGAGAAGCTGGGCCCACCCC |

**Table S3: List of oligonucleotides used for CRISPR/Cas9 experiments to target *NFKB2^D865G^***

| **Gene** | **Forward** | **Reverse** |
| --- | --- | --- |
| *NFKB2* | CCCTAACCATGACTCAGACCTCA | CCTCCCCTTCCCATGAGAATCC |
| *TBX19* | CCCCTGGACAAGGTGAGAGTT | GACTCCCGGGAATAATTGGCTTC |

**Table S4: List of primers used for PCR analyses, CAPS assay and Sanger Sequencing**

|  | **Time** | **Temperature** | **Cycle** |
| --- | --- | --- | --- |
| **Initial denaturation** | 1 min | 94°C |  |
| **Denaturation** | 10 sec | 98°C | 40 cycles |
| **Hybridization** | 30 sec | 62.5°C |  |
| **Elongation** | 30 sec | 68°C |  |
| **Final elongation** | 1 min | 68°C |  |

**Table S5: PCR process for *TBX19***

|  | **Time** | **Temperature** | **Cycle** |
| --- | --- | --- | --- |
| **Initial denaturation** | 1 min | 98°C |  |
| **Denaturation** | 10 sec | 98°C | 40 cycles |
| **Hybridization** | 10 sec | 66°C |  |
| **Elongation** | 30 sec | 68°C |  |
| **Final elongation** | 1 min | 68°C |  |

**Table S6: PCR process for *NFKB2***

| **Gene** | **Forward** | **Reverse** |
| --- | --- | --- |
| *ACTB* | CGGGAAATCGTGCGTGACATTAAG | GTAGTTTCGTGGATGCCACAGGA |
| *GAPDH* | CGGAGTCAACGGATTTGGTCGTAT | CAGCATCGCCCCACTTGATTTTG |
| *TUBB* | TGAGGGAAATCGTGCACATCCA | CCAAAAGGACCTGAGCGAACAGA |
| *BMP4* | ACCTCGGCCAAGTAACGGTAGT | TCCGGACTACATGCGGGATCTTTA |
| *FGF10* | TGTGCGGAGCTACAATCACCTT | CAGGATGCTGTACGGGCAGTT |
| *FGF8* | ACACCTTTGGAAGCAGAGTTCGAG | TGAAGACGCAGTCCTTGCCTTT |
| *HESX1* | CGCTCAGCTCGGGGAAAACAAA | ACCACGCTAGGGAATGAAATCCCA |
| *LHX3* | TGGTGCAGGTTTGGTTCCAGA | ATTTCCGCCAAGGAAGGCTCAT |
| *NEUROD4* | CCAGTGACCGAGAGTCTGGA | ACCTCATTTTGGGAGCCCAGA |
| *NR4A2* | AGGTTCCAGGCGAACCCTGACT | AGGTCTGCGAAGCCAGGGATCT |
| *PCSK1* | CAACTATGATCCAGAGGCTAGC | TCTGGTCCCGTGTTTGTTC |
| *PITX1* | GGCAACGTACGCACTTCACAA | TGCACTAGGCCGCTGAACT |
| *POMC* | CCCCTACAGGATGGAGCACTT | GATGGCGTTTTTGAACAGCGT |
| *PROP1* | GCAGTTGGAACAGCTGGAGTCA | AAGCAGTGGACTCTGGCAAGAA |
| *TBX19a* | TGAGAGCCAGCATGTGACCTAT | TGTGCATGTACGCAGAAGGGT |
| *TBX19b* | TGGCAGACGGATGTTTCCAGTC | CACCCATTCCCCGTTGACGTAC |
| *ZBTB20* | AACGGAAGAAACCCAAGACAGCT | AGCTTCAAAGTTCAGGCAGGGA |

**Table S7: List of primers used for qRT-PCR**

| **Antibody** | **Host species** | **Company and catalog number** | **Dilution** |
| --- | --- | --- | --- |
| Anti-NFKB2 | Rabbit | Sigma, HPA008422 | 1/500 |
| Anti-LHX3 | Mouse | DSHB, 67.4E12 | 1/500 |
| Anti-TBX19 | Rabbit | Sigma, HPA072686 | 1/300 |
| Anti-ACTH | Guinea Pig | NIDDK | 1/2000 |
| Anti-E-cadherin | Rat | Millipore, MABT26 | 1/500 |

**Table S8: List of primary antibodies used for immunostaining**
