## Additional detailed methods for "Modeling corticotroph deficiency with pituitary organoids supports the functional role of *NFKB2* in human pituitary differentiation"

**Method details**

**Culture and maintenance of hiPSC lines**

The 10742L healthy individual-derived hiPSC line was used in the study as the WT control (provided by the Cell Reprogramming and Differentiation Facility [MaSC], Marseille Medical Genetics, Marseille, France). Information about the hiPSC line is indicated in **Suppl. Table S1**. Two mutant lines carrying *TBX19^K146R/K146R^* and *NFKB2^D865G/D865G^* were generated using CRISPR/Cas9 editing from the control line. All hiPSC lines were cultured on six-well (Corning, #3335, New York, USA)-coated plates by Synthemax II-SC Substrate (working concentration at 0.025 mg/mL, Corning, New York, USA) and maintained undifferentiated in a chemically defined growth medium (StemMACs hiPSC-Brew XF human medium; MACS Miltenyi Biotec, Paris, France)^24^. hiPSC lines were maintained in a humidified incubator under conditions of 37°C, 5% CO_2_, with a daily change of medium, and passaged when cells reached 60-80 % confluency using enzyme-free ReleSR according to manufacturer’s recommendations (StemCell Technologies, Canada, #05872).

**CRISPR/Cas9 mediated genome editing of hiPSC**

CRISPR/Cas9 gene editing was used to create the *TBX19^K146R/K146R^* and the *NFKB2^D865G/D865G^* mutations using the method as previously described^25^.

The CRISPR/Cas9 was used to generate two mutant hiPSC lines from the 10742L hiPSC line:

1) The *TBX19^K146R/K146R^* line, harboring the c.437A>G, p.Lys146Arg *TBX19* (NM_005149) pathogenic missense variant.
2) 2) The *NFKB2^D865G/D865G^* line, harboring the c.2594A>G, p.Asp865Gly *NFKB2* (NM_001322934) pathogenic missense variant.

*Preparation of the CRISPR/Cas9 sgRNA plasmid*

For each targeted gene, sgRNA plasmid was prepared as previously described^26^. Briefly, we designed a sgRNA sequence within 20-nucleotides from the target site selected with the open-source CRISPOR tool (http://crispor.tefor.net/)^27^. This sgRNA was chosen specifically to guide the Cas9 enzyme to the targeted sequence, and Cas9 creates a double-strand break of DNA 3 base pairs upstream of protospacer-adjacent motif (PAM) sequence. The actual genome editing occurs in the process of repairing the double-stranded breaks created by the CRISPR-Cas9 system by using homology-directed repair (HDR) and non-homologous end joining (NHEJ).

The sgRNA sequence was used to generate *TBX19^K146R^* was as follow (5’>3’): AAGCTGACCAACAAGCTCAA (see also Table S2).

The sgRNA sequence used to generate *NFKB2^D865G^* was (5’>3’): GTGAAGGAAGACAGTGCGTA (see also Table S3).

We used the cAB03 (also known as pX459-pEF1alpha) vector was previously described^25^. Briefly, pSpCas9 (BB)-2A-Puro (PX459) V2.0 (Addgene plasmid # 62988, Addgene, Teddington, UK) was modified by the EF1 alpha promoter^25^.

The chosen sgRNA was cloned into the cAB03 open plasmid vector using the method previously described by Arnaud et al^25^. Briefly, oligonucleotide pairs were hybridized with a buffer containing Tris HCl (100 mM) and NaCl (500 mM) and placed in a thermocycler ramping from 95°C to 18°C at 0.05°C/s. cAB03 was opened with BbsI (New England Labs Inc., Ipswich, Massachusetts, USA), then ligated with annealed oligonucleotides by incubating with T4 DNA Ligase (New England Labs Inc.) 10 min at room temperature (RT). The ligation product was transformed in competent bacteria DH5α (Invitrogen, Waltham, Massachusetts, USA) following manufacturer’s instructions. After incubation for 1 h at 37°C, the bacteria were inoculated onto ampicillin-containing LB agar plates (100 μg/mL) and incubated overnight at 37°C. The next days, haft of the bacterial colony was replated onto an ampicillin LB agar plate and the other haft was extracted 10 min at 95°C in a lysis buffer containing 20 mM Tris HCL, 2 mM EDTA and 1% Triton 100X (pH8). Then, a PCR was performed to verify sgRNA insertion, using primer Pr1127 (5’-ACTATCATATGCTTACCGTAAC-3’) that binds to the backbone and the reverse oligonucleotide used to insert the sgRNA. The PCR product was put to migrate on a 2% gel to verify the insertion of the sgRNA (102 bp band) in the bacteria. Two positive clones were then sent for Sanger sequencing (Genewiz, Leipzig, Germany) to confirm the presence of the sgRNA in the plasmid after miniprep extraction (NucleoSpin®Plasmid kit, Macherey Nagel, Allentown, Pennsylvania, USA). A validated clone was then amplified, its DNA extracted by midiprep (NucleoBond®Xtra Midi EF kit, Macherey). The final vector expressed the corresponding sgRNA under the control of the U6 promoter, as well as *Cas9* under the control of the EF-1alpha promoter, and a puromycin resistance gene.

*Single-stranded oligo DNA nucleotides (ssODN) design*

We also designed donor sequences to generate *TBX19^K146R^* and *NFKB2^D865G^* missense mutations. A donor sequence is used for homology-directed repair, allowing the introduction (“knock-in”, KI) of single nucleotide variations, with low efficiency. To optimize the efficiency of KI editing, the donor sequences were designed as single-stranded oligo DNA nucleotides (ssODN), carrying the mutation of interest, and also included a silent (or blocking) mutation that mutate the PAM or disrupt the sgRNA binding region to prevent re-cutting by Cas9 after successful editing^28,29,25,30^. The ssODN contained 100 nucleotides with homology arms on each side of the target region. The ssODN sequence was as follows (see also **Suppl. Table S2 – Table S3)**:

For *TBX19^K146R^* (5’-3’): TGGATGAAAGCTCCCATCTCCTTCAGCAAAGTGAGGCTGACCAACAAGTTAAATGGAGGCGGGCAGGTACGAATGAGGCGGGCAGGCCTGGCCACCCGCT.

For *NFKB2^D865G^* (5’-3’):

TCCCATTCCTGTCCCCATTTACCCCCAGCAGAGGTGAAGGAAGGCAGTGCCTACGGGAGCCAGTCAGTGGAGCAGGAGGCAGAGAAGCTGGGCCCACCCC.

ssODN were synthesized by Integrated DNA Technologies (Coralville, Iowa, US) at Ultramer™ quality.

*Electroporation*

hiPSC were transfected with constructed sgRNA/Cas9 vectors by electroporation using the Neon Transfection System 100 µL kit (Invitrogen). A summary of the procedure is described in **Suppl. Figure S1**. After dissociation into single cells using Accutase® (Innovative Cell Technologies) for 4 min at 37°C, cells were centrifuged at 300 x g for 4 min. The cells pellet was washed with DPBS without Ca^2+^ and Mg^2+^. Cells were centrifuged one more time at 300 x g for 4 min. 1 x 10^6^ cells were seeded with 4 µg ssODN, 4 µg sgRNA/Cas9 vector, and 100 µL resuspension buffer R (Neon Transfection System 100 µL kit, Invitrogen). Cells were then electroporated with the Neon Transfection System using the following parameters: single pulse of 1100 V for 30 ms. The electroporated cell suspension was then flushed into a single well of a 6-well coated with Synthemax II-SC substrate as above containing StemMACS^TM^ iPS Brew XF medium supplemented with 10 µM of Y-27632. Of note, another well containing cells electroporated without plasmids was used as a control to define the best time window at which the puromycin treatment had to be stopped to select the electroporated cells. 24 hours after electroporation, the culture medium was replaced with StemMACS^TM^ iPS Brew XF medium supplemented with puromycin 1 µg/µL. After 24h of selection and checking for the complete death of all control electroporated cells, the puromycin-resistant cells were switched back to normal culture iPS-Brew medium and were maintained until clonal colonies could be observed.

*Clone isolation and screening*

On days 3 post-transfection, we used the cleaved amplified polymorphic sequences (CAPS) assay to estimate the efficacy of CRISPR/Cas9 in bulk transfected hiPSC populations to choose the number of colonies to be picked. CAPS analysis is one of the most widely used mutation detection methods^31^. A restriction enzyme recognition site is inserted in the donor template. CAPS primers and restriction enzymes were determined with the help of AmplifX software (version 2.0.0b3; https://inp.univ-amu.fr/en/amplifx-manage-test-and-design-your-primers-for-pcr). CAPS primers were listed in **Suppl. Table S4**. The CAPS products were separated and visualized by electrophoresis on a 2% agarose gel and analyzed for densitometry with ImageJ^25,30^.

After 10 days, the CAPS assay was also used to screen and identify the possible *knock-in* (*KI*) clones. Each individual colony arising from a single puromycin-resistant cell was manually picked under a microscope and cut in two halves: one half was used for genomic DNA extraction followed by CAPS screening and then confirmed by Sanger sequencing, the other haft was detached, gently fragmented and transferred to a single well of a Synthemax II-SC substrate coated 12-well plate containing pre-warmed StemMACS iPS-Brew XF medium supplemented with 10 µM Y-27632. These clones were maintained in culture until colonies were expanded enough to be passaged manually or using ReleSR. After expansion, each individual clone was then cryopreserved using StemMACS Cryo-Brew^32^ .

KI clones were screened by CAPS assay and confirmed by Sanger sequencing. For genomic DNA extraction, one haft of each of the colony was individually transferred into 0.2 mL PCR tube and centrifuged 5 min at 300 x g. After remove the supernatant, the pellet cells washed with 150 µL DPBS without Ca^2+^ and Mg^2+^ and centrifuged 5 min at 300 x g. The cells were re-suspended with a 20 µL mix containing: 1X of the 5X PrimeSTAR GXL Buffer (Clontech, Takara Bio, Shiga, Japan), proteinase K (0.167 mg/mL, Roche Diagnostics, Basel, Switzerland) and ddH_2_0. DNA extraction was performed using a thermocycler with the following settings: 3h at 55°C and 30 min at 95°C. Then, a 20 µL PCR reaction was performed using 1 µL of the extracted genomic DNA and 19 µL of a PCR mix containg 1X of the 5X PrimeSTAR GXL Buffer, PrimeSTAR GXL DNA polymerase (0.25 U/10 µL), dNTPs (160 µM each) and the specific primers for each genotypic situation reported in **Suppl. Table S4**. PCR reaction was performed using the PCR condition reported in **Suppl. Table S5 – Suppl. Table S6**. In the case of TBX19, the PCR product was digested with the MseI enzyme (New England Biolabs, NEB, Ipswich, Massachusetts, USA) whose recognition site is the blocking mutation site (there was not the restriction site at the target site). For the digestion 10 μL PCR product are combined with 0.5 μL of MseI, incubated at 37°C for 2 h or 0.05 μL of MseI overnight at 37°C. In the case of NFKB2, we used the BtsI enzyme (New England Biolabs, NEB, Ipswich, Massachusetts, USA) whose recognition site is at the target site. For the digestion 10 μL PCR product are combined with 0.5 μL of BtsI, incubated at 55°C for 2h or 0.05 μL overnight at 55°C. The samples were then deposited to migrate onto a 2% agarose gel and revealed to analysze the size of the amplicon for each individual clone.

The DNA of clones identified as possibly KI was purified using a DNA purification kit (Macherey-Nagel), and the samples were verified by the Sanger sequencing (Genewiz, Leipzig, Germany). Analyses of Sanger traces were done by Sequencher DNA sequence analysis software (version 5.4.6, Gene Codes corporation, Ann Arbor, MI USA, [http://www.genecodes.com](http://www.genecodes.com/)) to aligned sequences of KI clones with WT sequence to confirm the successful edition. Sanger sequencing primers are described in **Suppl. Table S4**. KI clones were kept, amplified and then cryopreserved.
