## Supplementary videos for "Modeling corticotroph deficiency with pituitary organoids supports the functional role of *NFKB2* in human pituitary differentiation"

### Slide 1
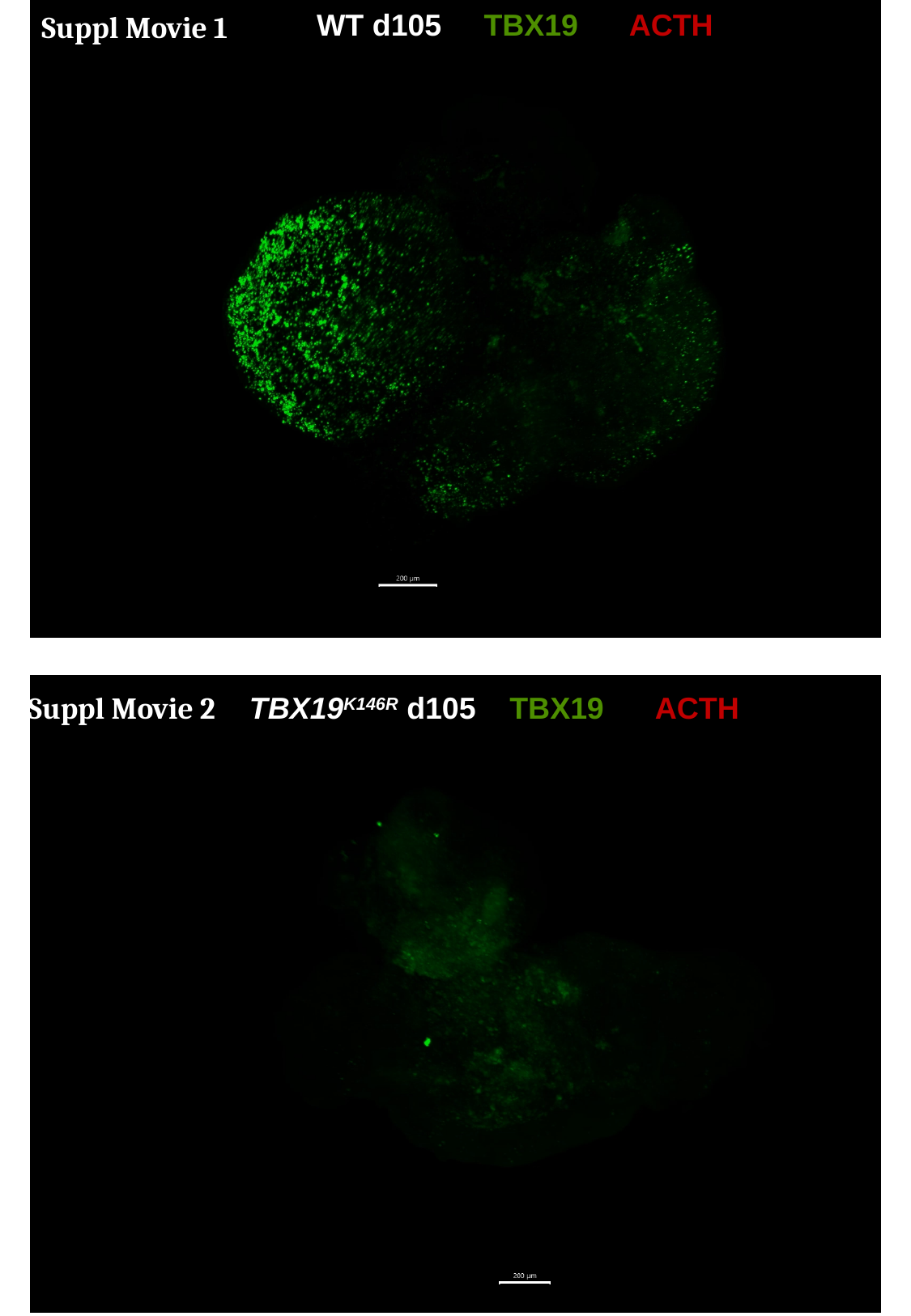

WT d105 ––TBX19 ACTH
Suppl Movie 1
Suppl Movie 2
TBX19^K146R^ d105 – TBX19 ACTH

### Slide 2
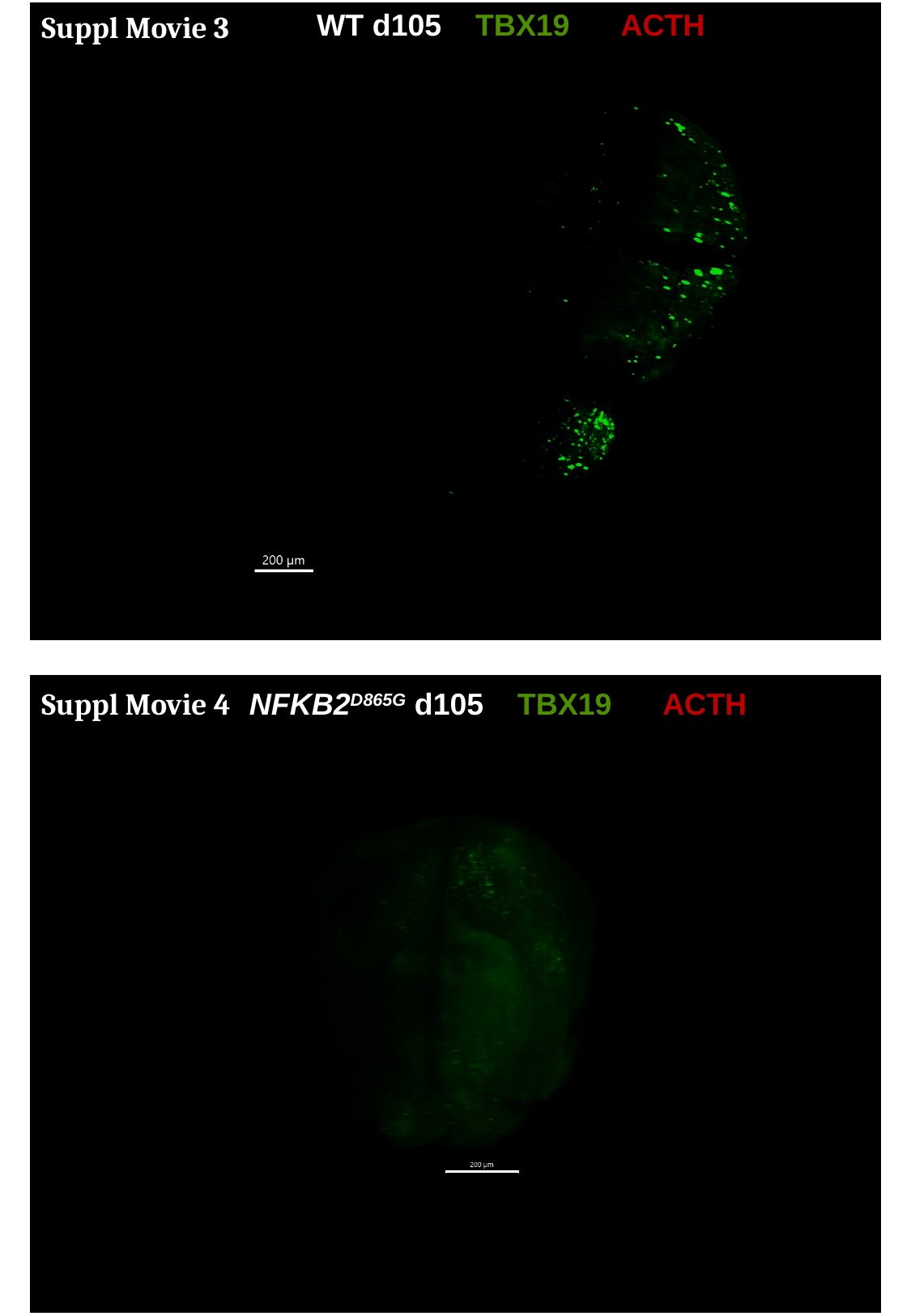

WT d105 – TBX19 ACTH
Suppl Movie 3
Suppl Movie 4
NFKB2^D865G^ d105 – TBX19 ACTH
